# Evidence for early kidney vasculature development by vasculogenesis and angiogenesis based on advanced in vivo chimeric model and depletion of CD31+/CD146+ intrinsic endothelial progenitors

**DOI:** 10.64898/2026.09.15.750731

**Authors:** Susanna Kaisto, Samar Ahmad, Sabrina Halmbacher, Virpi Glumoff, Laura Dönges, Irina Raykhel, Ilya Skovorodkin, Seppo J Vainio

**Affiliations:** Laboratory of Developmental Biology, Disease Networks Research Unit, Faculty of Biochemistry and Molecular Medicine, University of Oulu, Oulu, Finland; GeneCellNano, University of Eastern Finland, Kuopio, Finland; Institute of Biochemistry and Molecular Biology, Ulm University, Ulm, Germany; Research Unit of Biomedicine, University of Oulu, Oulu, Finland; The Unit of Measurement Technology (MITY), University of Oulu, Oulu, Finland; Kvantum Institute, University of Oulu, Oulu, Finland; Infotech Oulu, University of Oulu, Oulu, Finland

**Keywords:** Kidney, vascularization, xenotransplantation, CAM, CD146

## Abstract

Kidney diseases are a significant health concern worldwide, with a growing demand for fundamental treatments, such as generation of transplantable organs from programmed cells. The generation of kidney organoids has led to significant advances in disease modeling. However, failure to establish a robust endothelial vasculature network in kidney organoids remains a barrier to oxygen support and gas exchange in renal tissue engineering.

We targeted the development of early kidney vasculature system by advancing the classic mouse embryonic kidney organ culture and organoid model system by introducing a novel mini reservoir setup. The method improved integration with the highly vascularized chicken Chorion Allantois Membrane (CAM), enabling host blood flow to developing kidney organoids. We examined the origin and role of the CD31+ endothelial cells (ECs) and their CD146+ progenitors by applying a dissociated embryonic kidney approach. This enabled selected depletion of specific cell populations, followed by regeneration of the kidney organ primordia by reaggregation and placement on the CAM to assess organogenesis. The cell lineage and fate tracing with constitutively labelled cells revealed that the intrinsic endothelial CD146+ progenitor and those of CD31+ ECs in the kidney initiating organogenesis established the kidney vasculature. If the intrinsic EC progenitors were depleted, the mouse glomerular vascularization was rescued by the CAM derived ECs.

## INTRODUCTION

The kidney is a highly organized organ that performs several vital functions, including filtering blood constituents and concentrating urine while eliminating metabolic waste products. For these processes to become established, kidney vasculature assembly is essential. Progress has been made in identifying endothelial progenitor cells behind the kidney vasculature assembly and how such cells become integrated to construct a functional nephron during organogenesis. However, whether the kidney is vascularized by angiogenesis or by a combination of angiogenesis and vasculogenesis remains still a matter of debate (Stolz & Sims-Lucas, 2015; Munro & Davies, 2018; Cosma et al., 2024). Vasculature assembly mechanisms during ontogenesis are relevant not only to developmental biology but also to renal tissue engineering to respond to the high demand for kidney transplant therapies.

Several protocols have been developed to generate kidney organoids from mouse embryonic stem cells (mESC) and human induced pluripotent stem cells (iPSC) (Freedman et al., 2015; Takasato et al., 2016; Oxburgh et al., 2017; Li & Izpisua Belmonte, 2019; Nishinakamura, 2019; Koning et al., 2020; Raykhel et al., 2024). A significant limitation however for efficient progress is how to establish vasculature network with blood flow that would reach nephrons and integrate into developing and functional glomerular tufts (Freedman et al., 2015; Takasato et al., 2016).

The power of developmental model systems for vasculature analysis is their readiness for experimental manipulation of processes, availability of cell specific reporters and sophisticated tissue specific gene editing tools. These capacities offer critical tools to advance our understanding of kidney vascularization critical for development of functional kidney organoids towards better kidney disease modeling, drug screening, and tissue engineering. However, the evident current limitation is how to promote vascularization to support organoid growth in *ex vivo* settings (Halt et al., 2016; Hu et al., 2016; Munro et al., 2017; Daniel et al., 2018; Munro & Davies, 2018).

When mouse embryonic kidney is dissected and placed to culture, organogenesis and the associated nephrogenesis occur *ex vivo*. However, such kidney rudiments lack blood flow due to poor vasculature development and the flow generating source (Ryan et al., 2021). However, if embryonic kidney explants or organoids are transplanted under the kidney capsule, vascularization takes place rapidly as well as perfusion of a branched vascular network and maturation of kidney organoid development occurs (Sharmin et al., 2016; Bantounas et al., 2018; van den Berg et al., 2018; Raykhel et al., 2024). However, such implantation approach fails to offer easy access to real-time vascularization image analysis. Xenotransplantation to the chorioallantoic membrane (CAM) of the chick embryo would offer well accessible imaging coupled with tissue engineering in the context of rich vascular bead support (Garreta et al., 2019; Koning et al., 2022).

Earlier studies based on fluorescence-activated cell sorting (FACS)-mediated depletion of the CD31- and CD146-positive ECs suggested the presence of renal endothelial progenitors in mouse embryonic kidneys (Halt et al., 2016). These results were based in most part on dissected, *in vitro*-cultured, and experimentally induced nephrogenesis in embryonic kidney mesenchymal cells.

In the current work, we report an advanced xenotransplantation method using mini reservoirs that promote graft growth and vascularization. Using this novel method, we targeted the roles of the CD31+/CD146+ ECs in kidney vasculature development by taking use of the prior developed kidney rudiment cell dissociation and reaggregation (DisRe) technology (Junttila et al., 2015). The advanced CAM based mouse embryonic kidney grafting assay supported organogenesis and endothelial cell growth, differentiation, and blood flow to developing glomeruli and enabled more detailed insights into kidney vasculature assembly mechanisms in the chimeras.

## RESULTS

### Embryonic Chicken CAM mini reservoir culture technology enhances kidney organogenesis

The classical CAM grafting assay, where organ rudiments are cultured on a vasculature bed, has provided a useful platform to grow cells and embryonic organs (Ekblom et al., 1982; Sariola et al., 1983; Garreta et al., 2019). However, in the traditional CAM assay the placed organ rudiments often fail to attach to the CAM. Due to this reason, success to support organogenesis remains sub optimal (Tarnick & Davies, 2022). In line with the earlier studies, when we placed kidney rudiments dissected from E11.5 mouse embryos on CAM, graft survival rate remained around 62% (Fig. 1E) due to failure of the grafted kidneys to adhere to the CAM, as a result several organs floated on the CAM or dried during the first 24 h of culture and underwent necrosis.

**Fig. 1.**
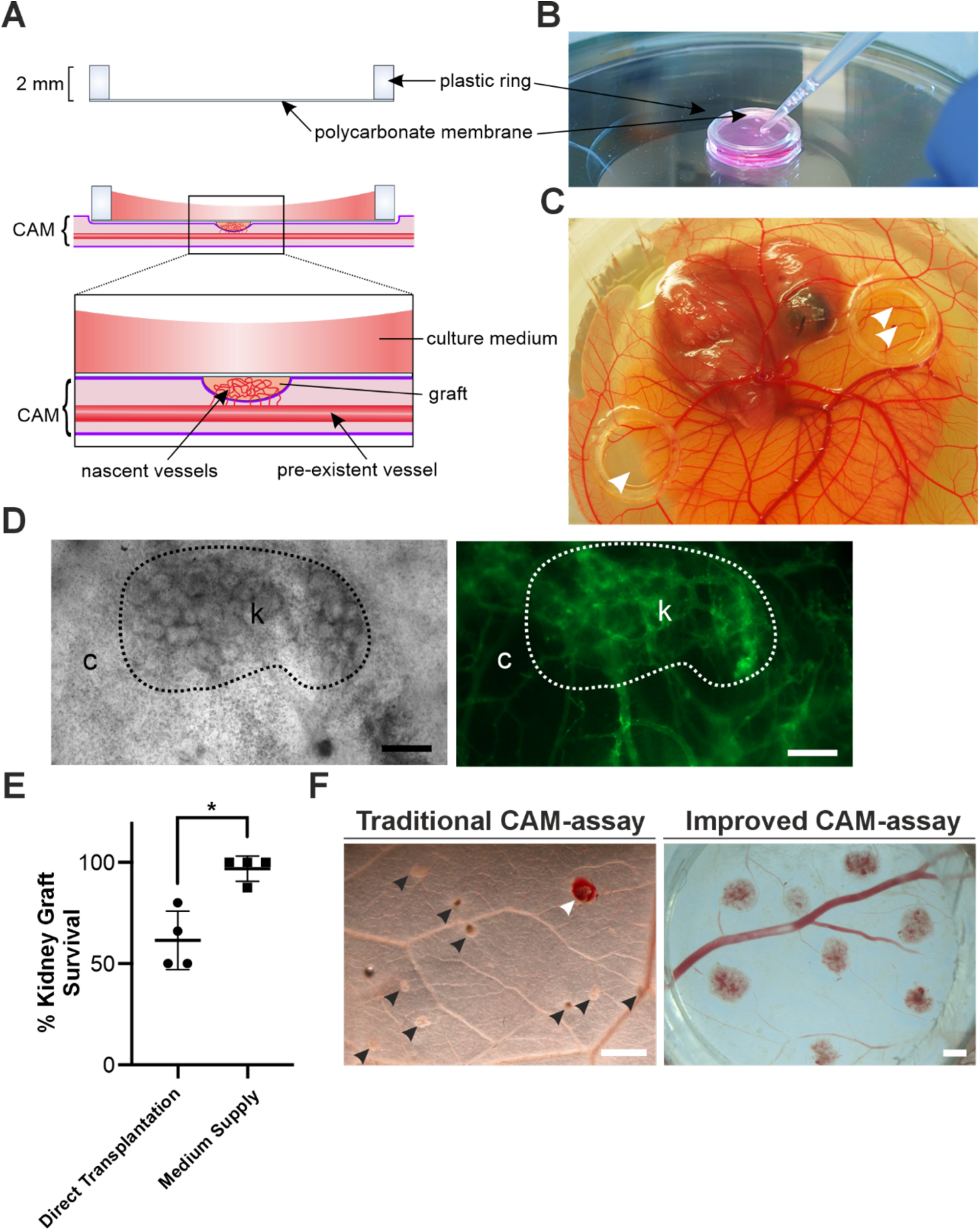
**A)** Mini reservoir-based CAM assay to culture kidney organ rudiments. A schematic picture of the applied mini reservoir. **B)** Mouse embryonic kidneys (E11.5) were placed on the membrane of inverted mini reservoirs, positioned on the CAM, and then supplied with culture medium in the well (**C).** In **C**) two mini reservoirs with E11.5 mouse embryonic kidneys (arrows) on the CAM of an HH41 stage chicken embryo after 5 days of cultivation are shown. **D)** Transmission light and fluorescence microscopic images of xenografted kidney (k) imaged on live chicken embryonic CAM (c) after injection of AF488-conjugated WGA lectin to the chicken blood circulation. Scale bar, 200 µm. **E)** The advanced culture assay increases survival of the E11.5 kidney grafts on the CAM. Percentage graft survival is plotted as mean ± SEM from n = 4 biological replicates. *P<0.05 using an unpaired t-test with Welch’s correction. **F)** The advanced CAM assay increases vascularization. Stereomicroscopic images of E11.5 kidneys grafted to *ex ovo* chicken CAM. In traditional method, 1 (white arrow) out of 10 (9 non-vascularized, black arrows), whereas in improved CAM assay, all 8 kidneys vascularized. Scale bar, 1000 µm.

The noted limitations in the classic CAM assay motivated us to further optimize the CAM organ/organoid culture assay. For this, we fabricated a custom mini reservoir by trimming commercial 12-well cell culture inserts, with 2 mm high polystyrene wall and outer diameter of 15.85 mm (Fig. 1A). The well bottom consisted of a 0.4 µm pore-sized transparent membrane. For the CAM assay, dissected kidneys were placed on the membrane of inverted mini reservoirs, which were then positioned on the CAM and filled with culture medium to prevent drying (Fig. 1B, 1C). This culture setup provided efficient media-based support for organogenesis both from the well and from the CAM provided nutrition. Moreover, the mini reservoir membrane compressed the kidneys and fixed them at the start of the culture, enabling efficient imaging of the organogenesis process. For instance, the developing vasculature system was labelled by intravenous injection of the fluorochrome-conjugated wheat germ agglutinin (WGA) in the CAM (Fig. 1D). The mini reservoir based-kidney rudiment CAM culture provided around 97% success to support organogenesis (Fig. 1E & 1F).

### Embryonic chicken CAM mini reservoir culture technology enhances kidney vasculature development

Given the need for vascularized organoid culturing technologies, a key aim was to develop an *ex vivo* method that would improve nephrogenesis and associated glomerulogenesis. The CAM supported vascularization is thought to be based on chimera formation between the chick and mouse cells (Koning et al., 2022). The chimeric vasculature network likely provides means for initiation of blood flow and the connected hemodynamic mechanical forces driven by the chick embryonic heartbeat pulsation.

To investigate putative impact of the CAM on kidney organogenesis, we assessed first the CD31+ endothelial network in intact E11.5 mouse kidneys transplanted for five days either on CAM or on Saxén-type organotypic cultures. Immunohistochemical staining with mouse-specific CD31 antibody depicted an extensive endothelial network in the kidneys grown on CAM compared to the classic culture setup (Fig. 2A). The murine CD31+ ECs had entered the glomerular structures in the CAM-grafted kidneys whereas the glomeruli on the Saxén-cultured kidneys lacked such populations (Fig. 2A). Thus, the CAM organ culture setup provided permissive conditions for ECs to develop renal vasculature.

**Fig. 2.**
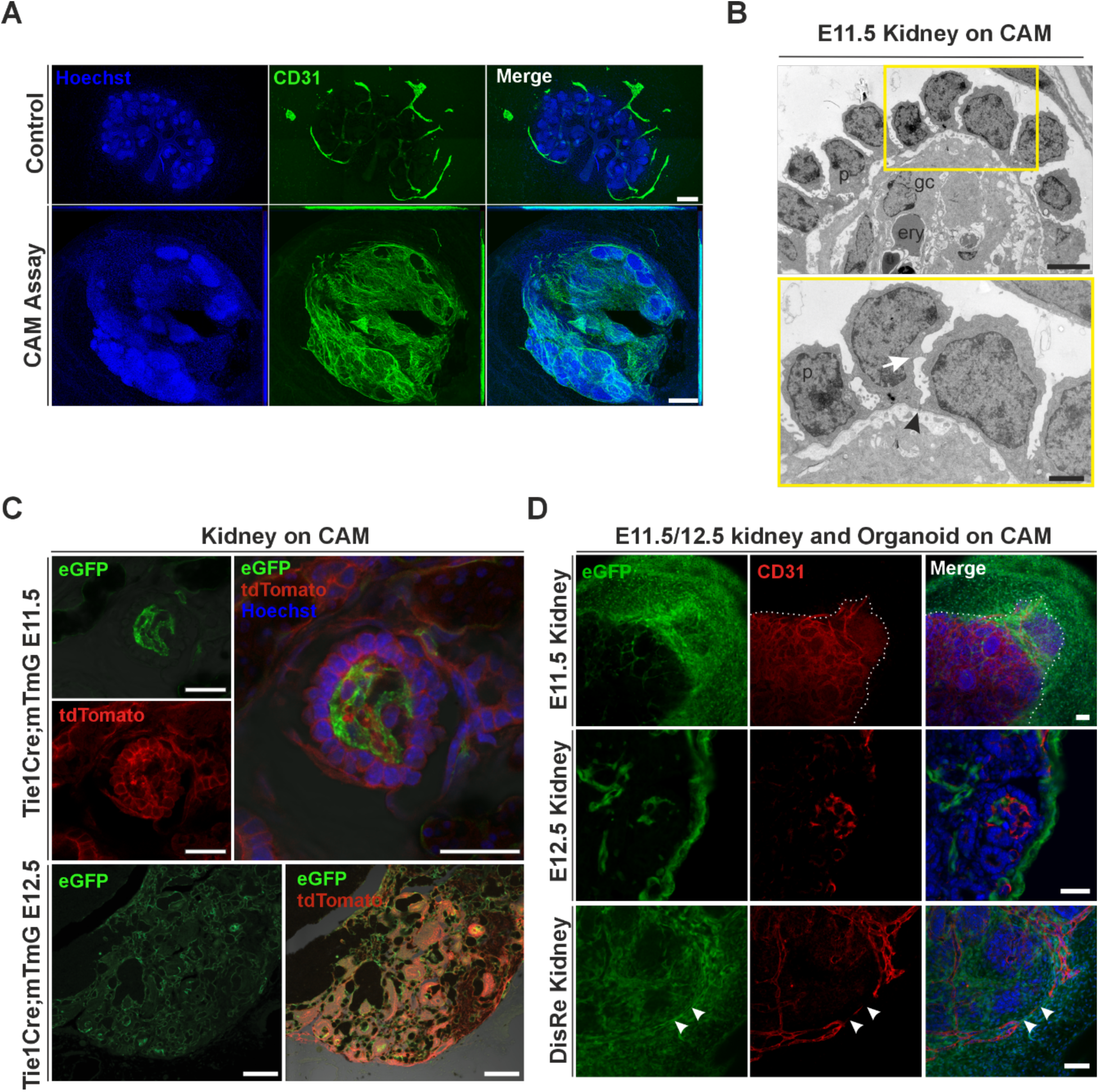
**A)** The advanced CAM assay promotes the survival and growth of the intrinsic endothelial cells (ECs). Embryonic kidneys at (E11.5) were dissected and cultured in classic organ culture setup (control) or as a xenotransplant on an eight-day-old embryonic chicken CAM. After five days, tissue samples were fixed and stained for endothelial marker CD31 (green). Scale bar, 200 µm. **B)** Transmission electron microscopy images of a glomerulus of an E11.5 kidney vascularized on CAM for seven days. Glomerular capillaries (gc), chicken erythrocytes (ery), podocytes (p), intercellular junctions (white arrow) and glomerular basement membrane (black arrow) are indicated. Scale bar, 10 µm (top) and 5 µm (bottom). **C)** ECs in vascularized xenografted kidney rudiments are mostly of donor origin. Tie1Cre;mT/mG mouse kidneys at E11.5 and E12.5 are xenografted to ten-day old chicken CAM and cultured for seven days. Glomerular tufts, that consist of eGFP+ mouse-derived ECs, can be observed. Scale bar, 20 µm (E11.5) and 100 µm (E12.5). **D)** Comparison of host versus graft origin of ECs in embryonic kidneys and kidney organoids placed on eGFP+ chicken embryonic CAM. Kidneys and kidney organoids at E11.5 and E12.5 were grafted to ten-day old eGFP+ chicken CAM and cultured for seven days. Samples were fixed and immunohistochemically stained with mouse-specific CD31 antibody and Hoechst as whole mounts (E11.5 kidney and DisRe organoids) or as cryosections (E12.5, middle panel). Arrows indicate chimeric blood vessels. Scale bars: 50 µm (whole mounts) and 20 µm (E12.5 cryosection).

Next, we analyzed by transmission electron microscopy (TEM) the degree of maturation of the renal corpuscles with the invaginated ECs (Fig. 2B). Primitive glomerular capillaries that had nucleated chicken erythrocytes, were observed in proximity of the podocyte-like cells. Junctions between these cells in proximity to the Glomerular Basement Membrane (GBM) were also noted (Fig. 2B). TEM images depicted fine structures in the developing glomeruli of embryonic kidneys grafted on CAM (Fig. 2B), in line with the prior reported data (Ichimura et al., 2017). Thus, the advanced CAM-based mini reservoir embryonic kidney culture supported glomerular vasculature development and podocyte-like cell foot process formation, thus representing critical steps for the functional maturation of glomeruli.

### Chicken embryonic CAM mini reservoir culture technology supports dual origin of the kidney endothelial progenitor cells

To determine the origin of the ECs in the mouse kidney rudiment, we used *Tie1^Cre^* transgenic mice (Gustafsson et al., 2001) in which Cre recombinase is targeted to the ECs. These mice were mated with a double fluorescent reporter mT/mG mouse (Muzumdar et al., 2007), where a membrane targeted tdTomato is expressed prior to Cre excision, whereas a membrane targeted EGFP is expressed post Cre excision, thus marking both recombined and non-recombined cells.

Embryonic kidneys were dissected from the Tie1^Cre^;mT/mG embryos at E11.5 or E12.5 and subjected to the mini reservoir-based CAM culture. Confocal images showed the presence of Tie1-expressing eGFP+ ECs throughout the kidney graft and often surrounded by chicken-derived erythrocytes (Fig. 2C) in the glomerular tufts, which mainly consisted of the eGFP+ mouse-derived ECs (Fig. 2C). We can conclude that mouse kidneys contain intrinsic ECs or/and EC precursors that when grafted on CAM become anastomosed with the chicken blood vessels and expand to form most of the renal and glomerular endothelium.

The Tie1^Cre^;mT/mG embryonic kidney approach only detects mouse-derived blood vessels. To track chicken embryonic CAM cells toward the mouse kidney rudiment ECs, we used Roslin Green chicken embryos ubiquitously expressing cytoplasmic EGFP. Mouse kidneys (E11.5/12.5) and organoids (E11.5) were xenotransplanted on the EGFP+ CAM. As noted with the Tie1^Cre^;mT/mG analysis, immunohistostaining revealed that the intrinsic murine CD31+ ECs had anastomosed with the chicken EGFP+ ECs (Fig 2D). Ingrowths of EGFP+ vasculature and CD31+/EGFP+ were also noted (Fig 2D), indicating chimeric vessels.

### Kidney organoids maintain nephrogenic potential upon depletion of CD31+ and CD146+ endothelial cells

To study the putative role of the intrinsic ECs and their presumptive progenitor cells in the kidney vascular development, we depleted CD31+ and/or CD146+ cells from dissociated E11.5 kidneys using immunostaining-based FACS (Fig. 3A, S1B). The remaining cells were reaggregated to form kidney organoids. Immunostaining of control kidney organoids revealed as expected extensive CD31+/CD146+ endothelial network, whereas CD31+/CD146+ single or double depleted lacked such network. Sporadic single CD31+ or CD146+ cells were noted in the reaggregates, indicating successful removal of EC precursor cells (Fig. 3B).

**Fig. 3.**
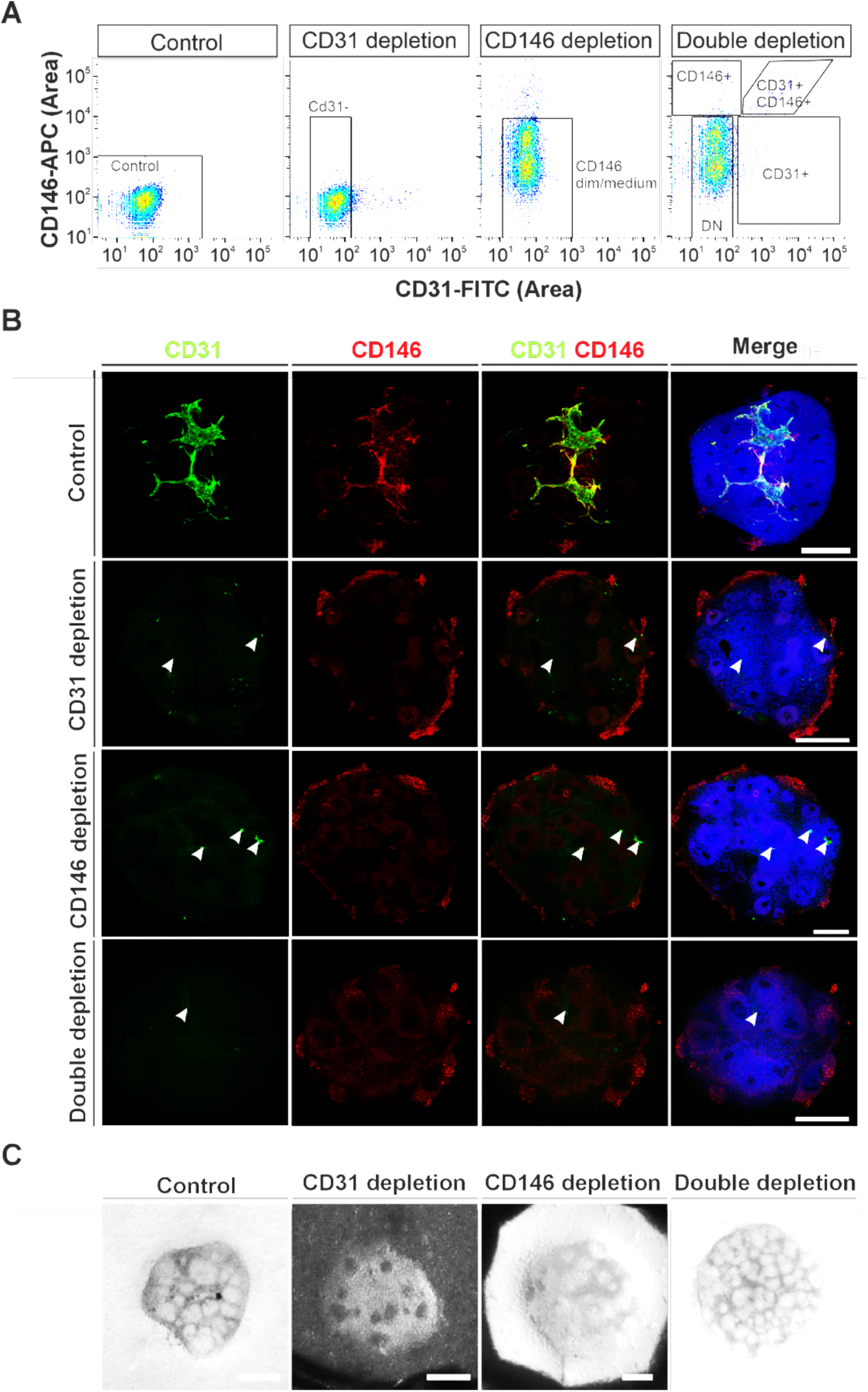
CD31 and CD146 depleted embryonic kidney organoids show nephrogenic potential. **A)** Dissociated wild-type (WT) murine kidneys at E11.5 were stained with CD31-FITC and CD146-APC antibodies separately or together, and subjected to FACS sorting of the CD146-, or CD31- or double-negative (DN) cells. Cells that were not immunostained served as controls. **B)** FACS leads to efficient depletion of the CD31+ and the CD146+ cells. FACS sorted kidney cells were reaggregated and cultured in the Saxén-type kidney culture for three days. The organoids were then fixed and immunostained as whole mount with CD31 and CD146 antibodies and Hoechst. Arrowheads indicate some remaining CD31+ cells in the kidney organoid. Scale bars: 200 µm. **C)** After three days of culture, stereomicroscopic images of organoids show nephrogenic potential in all FACS-sorted organoids. Scale bar: 200 μm.

The depletion of ECs was further confirmed through qPCR of the FACS-sorted CD31-/CD146-double-depleted cells (Fig. S2). The results showed a reduction in the expression of the CD31 and CD146, indicating efficient depletion, as well as other EC-specific genes (Tie1, Tie2, VEGFR2, and VE-cad). Angiogenesis-associated non-EC genes (VEGF, Angpt1 and Angpt2) and a UB epithelial cell marker (Gata3) were also reduced.

We next examined the nephrogenic potential of CD31, and CD146 single and double-depleted cells grown in organotypic culture setup. Sorted cells were reaggregated and the formed organoids were transferred to a Saxén-type organ culture for 3 days. Nephrogenesis was confirmed in all the organoids since renal structures were observed in stereomicroscopic images (Fig. 3C).

### CD31–/CD146– depleted kidney organoids are vascularized by chicken ECs when grafted to embryonic chicken CAM

When whole embryonic kidneys are grafted onto CAM, most blood vessels are donor-derived, with host ECs making some contribution to chimeric vessels and glomerular tufts (Kaisto et al., 2020; Oliveira et al., 2024). To determine whether intrinsic ECs are required to guide the chicken-derived vessels into the glomeruli, avascular kidney organoids were transplanted onto chicken CAM using the advanced mini reservoir setup (Fig. 4A). For this, E11.5 mouse kidneys were dissociated, stained with CD31-FITC and CD146-APC antibodies, and FACS-sorted to isolate CD31–/CD146– double-negative cells, which were reaggregated to form organoids. The organoids were precultured on mini reservoirs prior to transplanting on chicken CAM for five days (Fig. 4A). Dissociated kidney rudiments with CD31+/CD146+ cells still present were used for preparing control organoids (Fig. 4A).

**Fig. 4.**
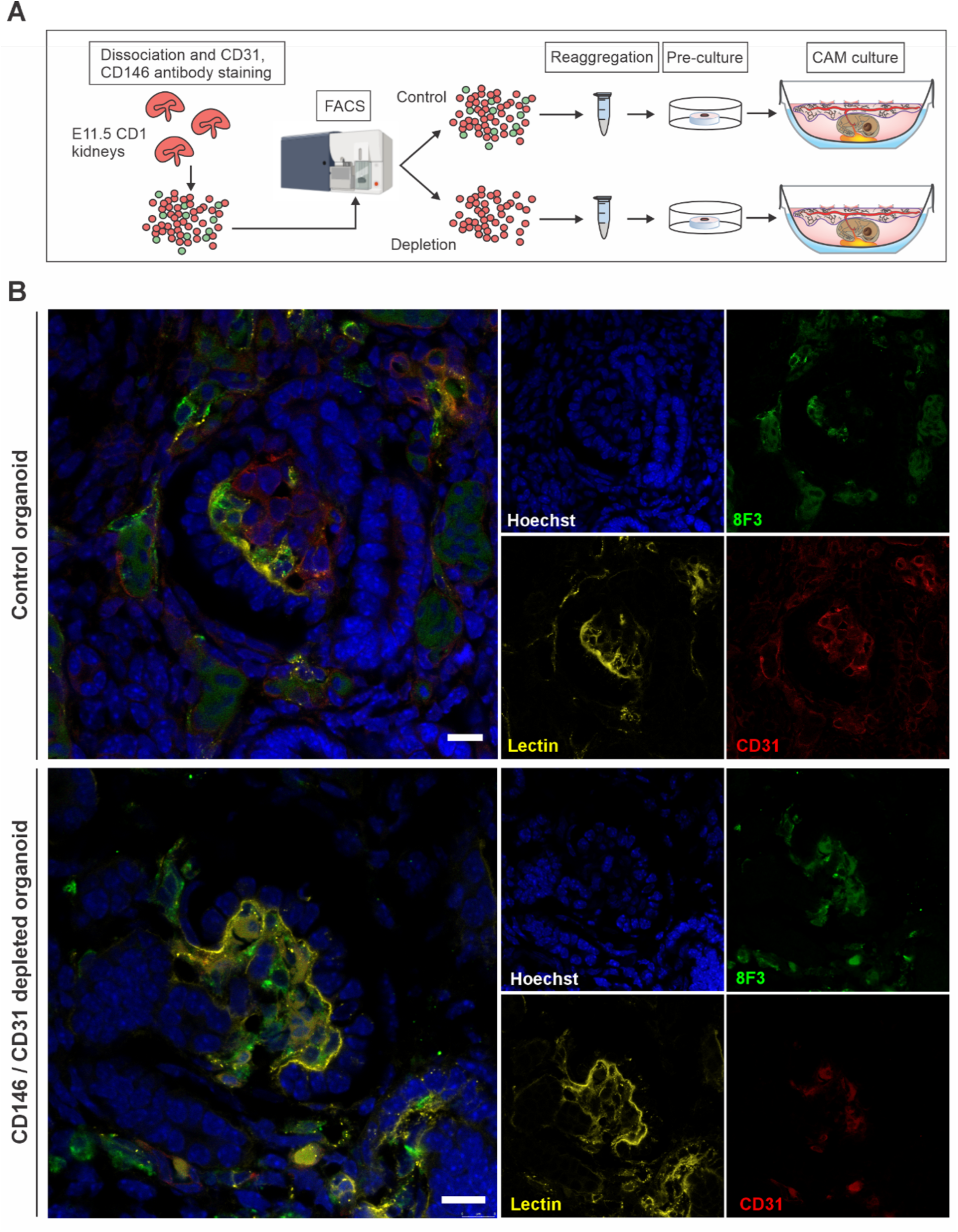
FACS-sorted kidney organoid grafted on CAM. **A)** Schematic picture of the experimental setup. **B)** Upon xenografting on CAM for 5 days, control kidney organoids were vascularized by both mouse-derived CD31+ ECs and host-derived chicken ECs, whereas CD31/CD146 double-negative kidney organoids were vascularized exclusively by chicken ECs. AF-conjugated WGA lectin, which was used to visualize perfused blood vessels, was injected into a CAM vein 20 min before organoid harvest and PFA fixation. A mouse-specific CD31 antibody was used to identify donor-derived ECs, and a pan-chicken antibody (8F3) was used to stain host-derived cells. Scale bars: 10 µm.

Immunohistochemical staining for mouse CD31 and chicken cells (using pan-chicken antibody 8F3) showed that both experimental and control organoids became vascularized when grafted on CAM. Moreover, WGA-AF546 stained vascular network was apparent in the grafted kidney organoids (Fig. 4B) as noted earlier in controls (Fig. 2). Notably both the CD31+ intrinsic mouse ECs and those of the 8F3+ CAM ECs had vascularized the developed glomeruli (Fig. 4B). In CD31-/CD146-depleted organoids, glomeruli only contained 8F3+ chicken ECs (Fig. 4B). Presence of occasional CD31+ cells outside of glomeruli in a few organoids likely reflects incomplete depletion or differentiation from residual progenitors (CD31-/CD146+). After five days on CAM (organoid day 7), glomeruli ranged from s-shaped body and comma-shaped body to glomerular tuft-stage.

## DISCUSSION

### Dual origin of embryonic kidney vascularization

The transplantation experiments indicate that the developing kidney, especially the renal vasculature, depends on systemic kidney related factors for their maturation. Embryonic kidneys and renal organoids are known to contain endothelial cell precursors, but the reasons why the maturation of renal blood vessels is limited in *in vitro* organotypic cultures (Bernstein et al., 1981; Sariola et al., 1983) but is supported when transplanted remains poorly understood. Two prominent factors absent in the traditional organ culture setup but present in the CAM culture assay could be the reason for this difference: 1) host-derived growth factors and signaling molecules, and 2) biomechanical effects from the blood flow.

We show that the intrinsic kidney primordia endothelial progenitor cells survive and develop into blood vessels when grown on the CAM. Such intrinsic renal ECs seem to form extensively the kidney associated blood vessels, however, with some contribution of chicken ECs (Fig. 2D). This implies that the vascularization of the embryonic mouse kidney occurs through a combination of vasculogenesis (intrinsic mouse ECs) and angiogenesis (chicken CAM vessels), which contradicts the earlier report in which blood vessels in grafted mouse kidneys were claimed to be of avian origin only (Sariola et al., 1983). Our data showed chicken-derived ECs in the developed glomeruli, which is in line with a previous study (Ekblom et al., 1982). Our results support the view that the metanephric kidney vascularizes through both by the vasculogenesis (Hyink et al., 1996; Robert et al., 1996; Loughna et al., 1997; Sims-Lucas et al., 2013) and the angiogenesis (Rogers & Hammerman, 2001; Dekel et al., 2003; Takeda et al., 2006; Munro et al., 2017) mechanisms.

### Role of the intrinsic kidney endothelial cells in kidney vascularization

We previously showed that endothelial progenitor cells in E11.5 murine kidney consist of a heterogenous pool of the CD31+ and the C146+ cells (Halt et al., 2016). We now obtained further evidence that the endogenous EC progenitors were essential for vascularization as depletion of CD31+ or CD146+ cells prevented the formation and development of a vascular network in the *in vitro*-cultured kidney organoid (Fig. 3B). Halt et al. (2016) reported the reappearance of CD31+ ECs in the CD31+ EC depleted kidney organoids (Halt et al., 2016). In the current study, upon depletion of CD146+ cells, a few CD31+ cells remained but were disorganized and no endothelial network formed (Fig. 3B), thus partially confirming the results of Halt et al., 2016. We also confirmed that removal of ECs does not affect nephrogenesis in renal organoids (Fig. 3C), however, the developing glomeruli remain avascular in this case.

### CD146+ cells as putative embryonic kidney endothelial cell progenitors

Halt et al. reported that the number of CD146+ cells decreased in mouse kidney between E11.5-12 (Halt et al., 2016). This and other observations led us to hypothesize that the CD146+ cells may represent progenitor cells for the renal ECs and go on to generate more mature CD31+/CD146– cells through a compound positive transition stage (Halt et al., 2016). Recently, iPSC-derived kidney organoids cultured on a microfluidic chip were shown to contain CD31+/CD146+ compound-positive cells predominantly, whereas organoids cultured on transwell inserts mostly contained CD31+ and CD146+ single positive ECs (Bas-Cristobal Menendez et al., 2022). These reports suggest that the CD31+/CD146+ compound-positive cells may indeed represent more mature ECs.

We found that the CD31-/CD146-double-negative cells exhibit reduced expression of EC marker genes including VEGFR2 in the depleted embryonic kidney organoids. The results are in line with the previous findings demonstrating that a majority of the CD31+ cells are also VEGFR2+ and Tie2+ (Murakami et al., 2019). Analysis of the kidney ECs with qPCR for the expression levels of other EC and EC progenitor marker genes, such as CD45, c-Kit, Sca, and Scl in the CD31+/CD146+ double-positive and the CD146 single-positive cells may be useful, since the CD146+ has been noted to serve as a co-receptor for the VEGFR2 (Jiang et al., 2012).

Since in the embryonic kidney rudiment majority of the CD31+ cells appear to be CD146+ (Fig. S1A), there is a possibility that the intrinsic ECs may originate from the CD146+ endothelial progenitor cells. The other possibility may be that the CD31+/CD146 + double-positive cells differentiate from a double-negative EC progenitor population.

Murakami et al. suggested the existence of a CD31–/VEGFR2–/Tie2– fraction in the E11.5 mouse kidney, possibly containing EC precursors that could contribute to the vascularization of transplanted kidney organoid more efficiently than the host ECs (Murakami et al., 2019). It would be interesting to test this hypothesis with CD146+/CD31– organoids grafted to chicken CAM.

### Kidney organoids depleted of intrinsic CD146+/CD31+ EC progenitors become vascularized by the CAM ECs

Previous reports on grafting mouse embryonic kidneys on CAM demonstrated predominant (if not complete) vascularization of the graft by host-derived ECs (Sariola et al., 1983; Sariola et al., 1984). We now provide evidence that the EC progenitors in the intact mouse embryonic kidneys and renal organoids in most part are represented by the recipient-derived cells. However, upon transplantation of the CD31/CD146 depleted organoids on the chicken CAM, kidney organoids became vascularized by the embryonic CAM endothelial cells. This key finding can now provide rationale for the discrepancy between our results and previously published findings. One interpretation would also be that endothelial cells may be sensitive and may indeed disappear from grafted kidneys before they would be vascularized by the CAM-derived endothelial cells. Our method of grafting on CAM with mini reservoirs provides now an optimized culture setup for survival of ECs in grafts and thus a useful assay for further cellular mechanistic studies.

## CONCLUSION

By using an improved chicken CAM assay, we studied the mechanism and origin of the renal endothelial cells during kidney vascularization. Our results underscore the critical role of blood flow in the formation of functional metanephric kidneys. Transplantation of embryonic kidneys to the chicken CAM offered permissive conditions for renal endothelial cell growth, enabling more advanced maturation than static organ culture. Both progenitor (CD146+) and fully differentiated (CD31+) endothelial cells present in E11.5 mouse kidneys contributed to renal vasculature formation, indicating that kidney vascularization happens through combined angiogenesis and vasculogenesis. Further studies are still needed to conclusively determine the source of the intrinsic kidney endothelial cells.

## Acknowledgements

We thank Paula Haipus, Hannele Härkman, Johanna Kekolahti-Liias and Ulla Saarela for technical assistance. We thank Veli-Pekka Ronkainen for confocal imaging support, Ilkka Miinalainen and Raija Sormunen for transmission electron microscopy support. We thank Sirpa Rannikko for FACS sorting, Andras Nagy for the GFP^+^ transgenic mice, and Erika Gustafsson for the *Tie1Cre* mice, Juha Partanen for the mT/mG mice. We thank Lauri Eklund for wheat germ agglutinate. We thank FACS core, light and electron microscopy facilities at the Biocenter Oulu, University of Oulu, Finland for support.

## Data availability

The data presented in this study are available in the article and supplementary material.

## Author contributions

Conceived and designed the experiments: SK, IS, SV. Performed the experiments: SK, IS, SH, VG, LD. Analyzed the data: SK, IS, SA, SH, VG. IR supervised LD for the quantification of kidney graft survival for the improved CAM assay method. Wrote the paper: SK, SA, IS, SV.

SV led the project, provided the infrastructure and finalized the writing of the paper. All authors approved the final version of the manuscript.

## Materials and methods

### Animals

Animal care and experimental procedures were in accordance with Finnish national legislation for the use of laboratory animals, the European convention for the protection of vertebrate animals used for experimental and other scientific purposes (ETS 123), and the EU Directive 2010/63/EU.

### Mouse models

All mice (*Mus musculus*) were maintained in the Oulu Laboratory Animal Center, housed in bedding environmental enrichments with unrestricted access to standard rodent chow and water. Wild-type (WT) embryonic kidneys were obtained from pregnant outbred CD1 (Charles River Laboratories) mice. *Tie1*^Cre^ (Gustafsson et al., 2001) mice expressing Cre-recombinase in the endothelial cells were bred with *tomato floxed Rosa26 GFP* (*mT/mG*) (Muzumdar et al., 2007) reporter mice to trace the mouse-derived endothelial cells in kidneys xenografted onto chicken CAM. B5/*EGFP* and *Tie1*^Cre^ mice were kept in the C57BL/6NCrl background (Charles River Laboratories). *mT/mG* mice were kept in C57BL/6NCrl background, but the mouse line was also backcrossed and kept in CD1 background. The appearance of the vaginal plug was considered to represent E0.5.

### Chicken embryos

Wild-type chicken embryos (*Gallus gallus domesticus*) were used in most of the xenotransplantation experiments. Fertilized Hy-Line White (Hy-Line international) and Nick Chick White (H&N International) chicken eggs were obtained from Haaviston Siitoskanala, Panelia, Finland. “Roslin Green” transgenic chicken embryos with ubiquitous cell expression of cytoplasmic eGFP (McGrew et al., 2008) were used to trace chicken endothelial cells in the CAM assay. Fertilized Roslin Green chicken eggs were obtained from National Avian Research Facility, Edinburg, UK.

### Preparation of chicken embryos: *Ex ovo* cultures

Upon arrival, the fertilized eggs were stored in a refrigerator at +12°C for a maximum of 14 days. To reinitiate the development, the eggs were placed in a 37.5°C humified egg incubator (Grumbach GmbH) on an automatic egg turner (OLBA B.V.) to reach the Hamburger-Hamilton developmental stages HH 17-18. Before opening the egg, the shell was wiped with a paper towel sprayed with 70% ethanol solution. The egg was opened from underneath by making a vertical crack in the shell on the dorsal side of the embryo. The entire content of the egg was transferred to sterile glass bowls, around 70% filled with sterile, prewarmed (37°C) water covered with cling film that was touching the water so that a rounded bed was formed for the egg; the method modified from (Schomann et al., 2013). The embryos were covered with a sterile glass or plastic petri dish before being placed in a 37°C cell culture incubator (90% humidity, 5% CO2). The embryos were allowed to develop for an additional 6 – 7 days until the CAM was developed enough for the CAM assay to be performed.

### Dissection and organ culture

Murine embryonic kidneys were isolated using previously described methods (Halt & Vainio, 2012). Whole kidney explants and kidney organoids were cultured at 37°C, and 5% CO_2_ in Dulbecco’s modified Eagle’s medium (DMEM)+ GlutamaxTM (5.56 mM glucose, 1 mM pyruvate) (Gibco) supplemented with 10% fetal bovine serum (FBS) (HyClone) and 0.5% penicillin-streptomycin (Sigma). The organ culture system used was a Saxén-type organ culture system (Saxén & Toivonen, 1962), where the kidneys and explants are cultured in the air-liquid interface on a polycarbonate filter (pore size 1.0 µm; Whatman) supported by a metal grid.

### Dissociation of kidneys into single cells

Murine embryonic kidneys were dissociated by incubating them for 30 – 45 min in 40 µl collagenase IV (Worthington) and 5 µl DNase I (New England Bio Labs) diluted in 280 µl of phosphate-buffered saline (PBS) at 37°C and disrupting the tissue by pipetting every 5 min. After the dissociation, the enzymatic reaction was inactivated by adding 1000 µl DMEM, 10% FBS, and 0.5% penicillin-streptomycin. The cells were centrifuged for 4 min at 1380 relative centrifugal force (RCF), and the pellet was resuspended in 100 µl PBS with 0.5% FBS.

### Fluorescence-activated cell sorting

Three different antibody labeling were performed. The dissociated mouse embryonic kidney cells suspended in PBS supplemented with 0.5% FBS were labeled with anti-CD31-fluorescein (FITC) eBioscience™ (Thermo Fisher Scientific Cat# 11-0311-82, RRID: AB_465012) and anti-CD146-allophycocyanin (APC) REAfinity™ (Miltenyi Biotec Cat# 130-118-408, RRID: AB_2751508). For single antibody staining, CD31-FITC was diluted 1:100 and incubated for 30 min on ice, CD146-APC was diluted 1:20 and incubated for 10 min at +4°C and when double staining, the incubation was first 20 min with CD31-FITC on ice and after addition of CD146-APC the incubation was continued with 10 min at 4°C. The cells were incubated while protected from light. The total volume was 100 µl in each case. All centrifugation steps were performed at 1380 RCF for 4 min. The cells were centrifuged and washed twice with 500 µl PBS with 0.5% FBS, then centrifuged and resuspended in 300 µl FluoroBrite^TM^ DMEM (Gibco™, Thermo Fisher Scientific Inc.). One drop of NucBlue™ Fixed Cell ReadyProbes™ Reagent 4′,6-diamidino-2-phenylindole (DAPI) (Invitrogen^TM^, ThermoFisher) was added immediately before flow cytometry. Cells were sorted using FACS Aria IIIu flow cytometer and collected in LoBind Eppendorf tubes (Eppendorf) containing 300 µl of kidney culture medium. The gating strategy of CD31-FITC against CD146-APC and the cell populations chosen for sorting are shown in Fig. S1B. Cell debris and dead cells were excluded from the analysis based on scatter signals and DAPI fluorescence. Flow cytometry data were analyzed with FlowJo^TM^ v10 Software (BD Biosciences).

### Embryonic kidney aggregates

The FACS-sorted cells were centrifuged for 4 min at 1380 RCF, and the cells for kidney organoid formation were resuspended in complete medium containing 10 µmol / l Y27632 (Enzo Life Sciences). The cell suspension was divided into 1.5 ml LoBind Eppendorf tubes, each containing a minimum of 5×10^4^ cells on average and centrifuged for 20 min at 1380 RCF and incubated at 37°C for 24 h. After this step, the culture medium was replaced with a standard kidney culture medium.

### Preparation of mini reservoirs and CAM-assay

The preparation of the modified cell culture inserts “mini reservoirs” is described elsewhere (Kaisto et al., 2020). Briefly, the membrane rings used in the improved CAM assay were modified Thincert^TM^ 12-well cell culture inserts (Greiner Bio-One) with 0.4 µm-pore polyethylene terephthalate (PET) membrane. The inserts’ sides were cut down to a height of 2–3 mm using a rotary multitool with a circular saw blade or a handsaw. The upper rim was then polished with a scalpel to remove sharp edges. The modified inserts were sterilized in 70% ethanol, washed in autoclaved double-distilled water, and dried in a laminar hood. Prior to culture, the mini reservoirs were rinsed in PBS and a complete culture medium (high-glucose DMEM–10% FBS–1% penicillin-streptomycin). The mini reservoir was placed on a petri dish on top of a small amount (400 µl) of complete medium with membrane facing up. One or several embryonic mouse kidneys or kidney organoids were arranged on each membrane with a pipette or a glass capillary tube. The samples were let to attach for 2–24 h in a cell culture incubator before transferring the cultures on day 9–10 chicken CAM. When placed on the CAM, the mini reservoir was positioned with the membrane facing the CAM, allowing the membrane to cover the grafts and hold them in place. The mini reservoirs were placed in the periphery of the CAM so that they would not disturb the embryo. A 300 µl aliquot culture medium was added inside the ring. CAM assays were kept for a maximum of 7 days, and fresh culture medium was added to the chamber every other day.

### Visualizing blood vessels and blood flow in CAM cultures

Fluorescent lectins were used to visualize perfused blood vessels. A total of 50 µl of 1 mg/ml Alexa Fluor® AF488- or AF549-conjugated wheat germ agglutinin (WGA) lectin (Invitrogen) together with 5 µl Fast Green solution (Sigma) was injected into a CAM vein in the host chicken embryo on the last day of the CAM assay. The chicken was euthanized after 20 min, and the kidney graft was collected and fixed with 4% paraformaldehyde (PFA).

### Quantitative real-time PCR (qRT-PCR)

Total RNA from cells, embryonic kidneys, or kidney organoids was extracted with a RNeasy kit (Qiagen) according to the manufacturer’s recommendations. Complementary DNA (cDNA) synthesis (First Strand cDNA Synthesis Kit, Thermo Fisher) was performed following standard protocols. qPCR analyses were displayed with SYBR Green (Agilent) by a CFX96 Real-Time PCR machine (Bio-Rad). The Brilliant III Ultra-Fast SYBR® Green QPCR Master Mix (Stratagene/Agilent Technologies) was used according to the manufacturer’s instructions. The qPCR cycling conditions were as follows: initial denaturation at +95°C for 3 min, followed by 40 cycles of +95°C for 5 sec and +60°C for 10 sec, and finally performing a melt curve at +65 – 95°C with an increment of 0.5°C for 5 sec. All primer sequences are given in Table S1. The validity of the primers was assumed for the primers published elsewhere, and for the primers we designed, primer validation was performed. The qPCR experiments were performed as three biological triplicates using mouse *Gapdh* gene as a reference gene. For the comparison of the gene expression between CD146^-^/CD31^-^ cells and control cells, the fold-change was calculated by the ΔΔCt method (*relative fold change = 2^-ΔΔCT^*).

### Immunohistochemistry

CAM-cultured kidneys and kidney organoids were collected from the CAM assay and washed in ice-cold PBS before PFA treatment. The PFA fixation step was from 10 min to 24 h, depending on the sample size. After fixation, samples were washed twice in PBS and stored at +4°C. For the cryosectioning, the explants were incubated first in 15% sucrose/PBS, followed by 30% sucrose/PBS, and a 1/1 mixture of 30% sucrose/PBS, and finally in the optimal cutting temperature compound (Tissue-Tek, Sakura Finetek) for a minimum of 30 min in each solution, and then mounted in plastic containers in which they were frozen in liquid nitrogen. The sections (5 mm cut with Leica Cryotome CM3050S or CM1950; Leica, Biosystems) were dried for 1 h at RT before staining. For immunostaining, the samples were blocked for 1 h in 0.1% Triton-X100/ 1% BSA/ 10% goat serum/ 0.2 M glycine-PBS at RT. Incubation of the samples with primary antibodies was performed in PBS-glycine or blocking buffer. After staining overnight at +4°C (or RT for nephrin) with primary antibodies, the samples were washed six times for 30 min with PBS. The samples were then incubated with the respective secondary antibodies and Hoechst (#62249, ThermoFisher Scientific) for 2 h at RT. Fluorescently conjugated primary antibodies were incubated together with secondary antibodies. The samples were mounted onto glass slides in a mounting medium (Immumount, ThermoScientific^TM^). The antibodies used for immunohistochemical staining in this study are listed in Table S2.

### Light microscopy

Bright-field and fluorescent images of xenotransplanted kidneys were taken with an Olympus SZ40 stereo microscope with an Olympus OM-D E-M10 Mark II camera. Fluorescent imaging of live chicken CAM injected with fluorescent lectin was performed using a Zeiss Axio Zoom.V16 fluorescence stereo microscope. Confocal imaging was performed using a Zeiss LSM 780 microscope (Zeiss) with Zen software or Leica SP8 FALCON confocal with a DMI8 microscope using LAS X 3.5.2 acquisition software. Subsequent data analysis was carried out with Zeiss Zen software. A maximum-intensity projection was performed to visualize endothelial cells. Only one layer of the Hoechst scans was selected for the presentation.

### Electron microscopy

Kidneys and organoids chosen for transmission electron microscopy (TEM) were fixed in 1% glutaraldehyde and 4% formaldehyde in 0.1 M phosphate buffer, postfixed in 1% osmium tetroxide, dehydrated in acetone, and embedded in LX-112 (Ladd Research Industries). The samples were processed and analyzed by the Biocenter Oulu electron microscopy (EM) core facility. Samples were examined using the Tecnai G2 Spirit transmission electron microscope (FEI), and images were captured with a charge-coupled device camera (Quemesa, Olympus Soft Imaging Solutions GMBH).

### Statistical analysis

Results are expressed as means ± standard deviation with a minimum of n = 3. Quantification of the kidney graft survival was performed using an unpaired t-test with Welch’s correction. Mean differences in mRNA expression were analyzed by one-way ANOVA followed by Tukey’s *post hoc* test. To ensure that the data were normally distributed, Levene’s test of homogeneity of variance was applied. A Welch test was additionally conducted when homogeneity of variances was not given. P-value < 0.05 was considered significant. Prism 9 (GraphPad) or SPSS Statistics (IBM) were used for statistical testing and graph preparation.

