## Supplementary Files for "Evidence for early kidney vasculature development by vasculogenesis and angiogenesis based on advanced in vivo chimeric model and depletion of CD31+/CD146+ intrinsic endothelial progenitors"

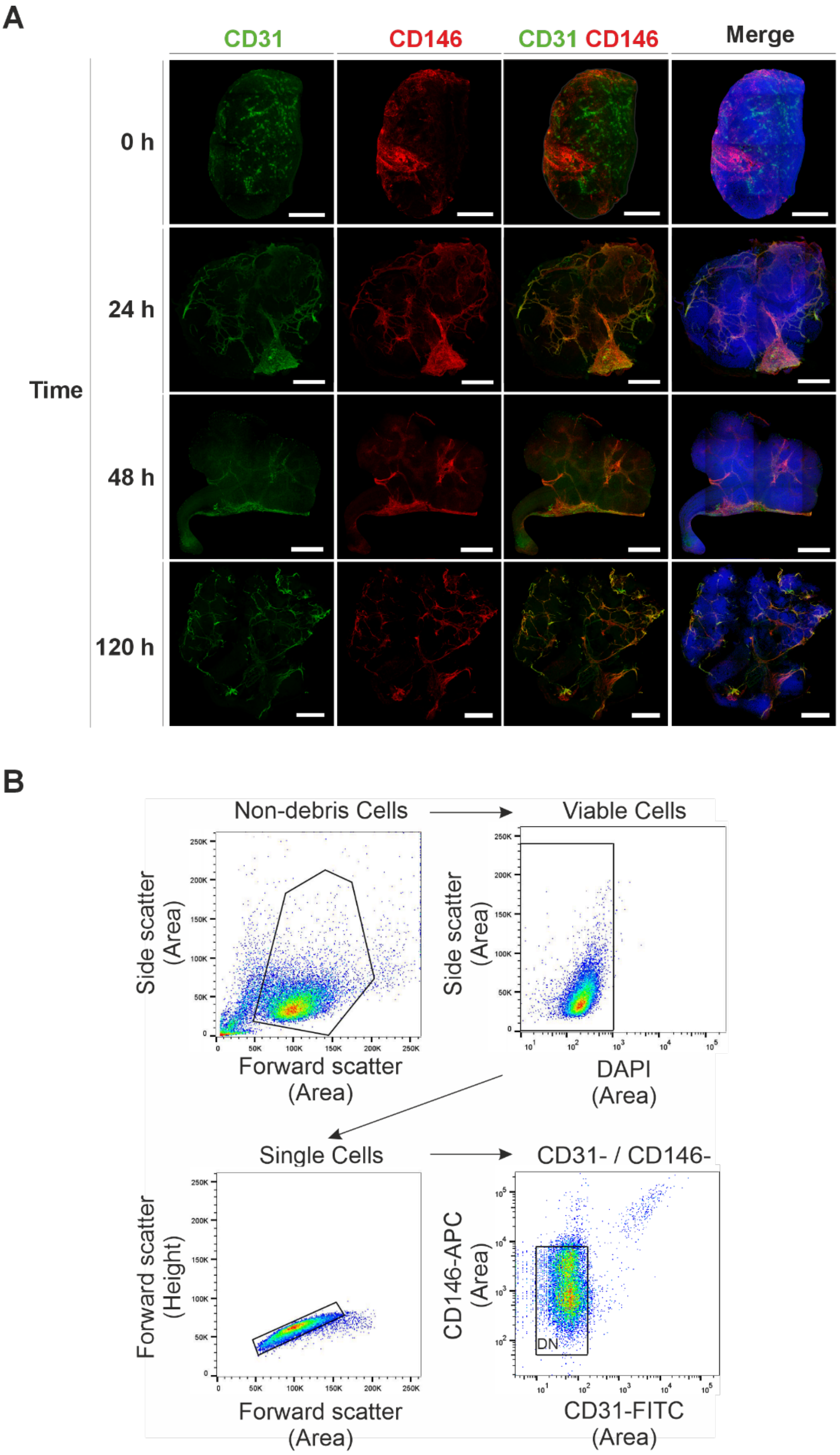
A
CD31
CD146
CD31 CD146
Merge
0 h
24 h
48 h
120 h
Time
B
Non-debris Cells
Viable Cells
Single Cells
CD31- / CD146-
Side scatter (Area)
Forward scatter (Area)
DAPI (Area)
Forward scatter (Height)
CD146-APC (Area)
CD31-FITC (Area)
DN

**Suppl.Fig.1**

**Fig. S1. A)** Distribution of CD31+ and CD146+ cells in primary embryonic kidneys grown on organotypic kidney culture. E11.5 mouse embryonic kidneys were dissected and either fixed immediately (0 h) or cultured and fixed after 24 h, 48 h, or 120 h. Whole mount immunohistochemical staining was performed with CD31, and CD146 antibodies and nuclei were stained with Hoechst. Scale bars: 200 µm (0 h, 24 h, 48 h), and 300 µm (120 h). **B)** ACS gating strategy of endothelial cells and their precursor cells based on CD31 and CD146. Representative FACS plots for the gating strategy to sort live CD31– and CD146– cells. E11.5 kidneys were pooled, dissociated, and double-labeled with CD31-FITC and CD146-APC antibodies. Cell debris and dead cells were excluded from the analysis based on scatter signals and 4',6-diamidino-2-phenylindole (DAPI) fluorescence. Flow cytometry data were analyzed with FlowJoTM v10 Software (BD Biosciences).

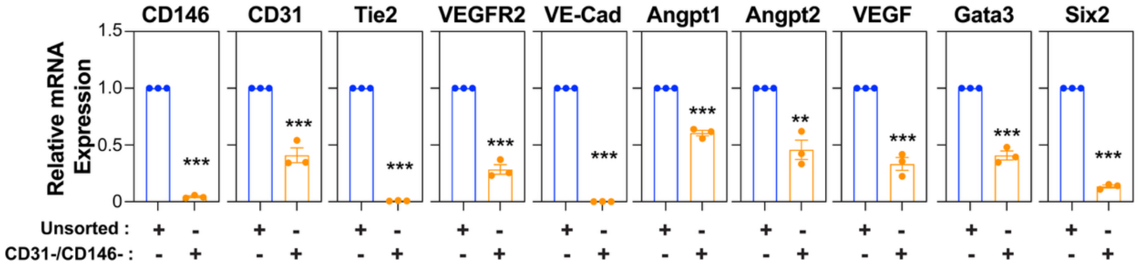


**Suppl.Fig. 2**

**Fig. S2.** Endothelial cell depletion reduces the expression of endothelial cell marker genes and angiogenesis-related genes. Total RNA was isolated from the FACS-sorted CD31$^{-}$/CD146$^{-}$ cell population and unsorted dissociated E11.5 mouse kidney cells and used to prepare cDNA and perform PCR reaction. Data is presented as mean ± SEM (n=3) of relative expression levels in FACS-sorted CD31$^{-}$/CD146$^{-}$ double negative cells with respect to unsorted dissociated E11.5 mouse kidney control cells normalized to 1. Statistical significance: *P<0.05; **P<0.001; ***P<0.001 using unpaired t-test.

**Table S1: Primers for gene expression analysis**

| qPCR Primers | | |
|---|---|---|
| **Gene** | **Forward Primer** | **Reverse Primer** |
| Angpt1 | CATTCTTCGCTGCCATTCTG | GCACATTGCCCATGTTGAATC |
| Angpt2 | TTAGCACAAAGGATTCGGACAAT | TTTTGTGGGTAGTACTGTCCATTCA |
| CD146 | AGGACCTTGAGTTTGAGTGG | CAGTGGTTTGGCTGGAGT |
| CD31 | GAGCCCAATCACGTTTCAGTTT | TCCTTCCTGCTTCTTGCTAGCT |
| VEGFR2 | GCCCTGCTGTGGTCTCACTAC | CAAAGCATTGCCCATTCGAT |
| Gapdh | CCAGCAAGGACACTGAGCAA | GGGATGGAAATTGTGAGGGA |
| Gata3 | CTCGGCCATTCGTACATGGAA | GGATACCTCTGCACCGTAGC |
| Six2 | CCGCGAGCTCACAAAATCC | CTTCTCCGCCTCGATGTAGT |
| Tie2 | ATGTGGAAGTCGAGAGGCGAT | CGAATAGCCATCCACTATTGTCC |
| VE-cadherin | TCCTCTGCATCCTCACTATCACA | GTAAGTGACCAACTGCTCGTGAAT |
| VEGF | GGAGATCCTTCGAGGAGCACTT | GGCGATTTAGCAGCAGATATAAGAA |

**Table S2: List of antibodies**

| **Reagent or Resource** | **Source** | **Identifier** |
|---|---|---|
| **Antibodies** | | |
| CD31 | BD Biosciences | 550274 |
| CD31 | Abcam | ab28364 |
| CD31-FITC | Thermo Fisher Scientific | 11-0311-82 |
| CD146-APC | Miltenyi Biotech | 130-118-408 |
| AF488-conjugated Goat anti-mouse IgG | Thermo Fisher Scientific | A-11001 |
| AF488-conjugated goat anti-rabbit IgG | Thermo Fisher Scientific | A-11008 |
| AF488-conjugated goat anti-rat IgG | Thermo Fisher Scientific | A-11006 |
| AF546-conjugated Goat anti-rabbit IgG | Thermo Fisher Scientific | A-11010 |
| AF546-conjugated goat anti-rat IgG | Thermo Fisher Scientific | A-11081 |
| AF647-conjugated goat antimouse IgG | Thermo Fisher Scientific | A-21235 |
| AF647-conjugated goat antirabbit IgG | Thermo Fisher Scientific | A-21244 |
| AF647-conjugated goat anti-rat IgG | Thermo Fisher Scientific | A-21247 |
